# The effects of host phylogenetic coverage and congruence metric on Monte Carlo-based null models of phylosymbiosis

**DOI:** 10.1101/2025.02.07.637028

**Authors:** James G. DuBose

## Abstract

Variation in host-associated microbial communities often parallels patterns of phylogenetic divergence between hosts, a pattern known as phylosymbiosis. Understanding of this phenomenon relies initially on quantifying phylosymbiotic signals from across a broad range of host taxa. Quantifying signals of phylosymbiosis is typically achieved by calculating how congruent a host’s phylogenetic tree is with a dendrogram that represents patterns of dissimilarity in their associated microbial communities. To statistically assess the degree of congruence, several studies have constructed null models using a Monte Carlo approach to randomly sample trees. Although this approach is becoming more common, it has several features that warrant benchmarking to advise its further use. This approach relies on quantification of congruence between a host’s phylogenetic tree its microbial community dendrogram. Therefore, it is important to establish how choice of congruence metric influences null model-based inferences. Furthermore, phylosymbiotic signals may manifest at different scales of host divergence, and it is important to establish the extent of host phylogenetic breadth needed to reliably detect a phylosymbiotic signal. To help improve our study of phylosymbiosis, here I examine how power and type 1 error (false positive) rates associated with this approach varies with choice of congruence metric and host phylogenetic coverage. Furthermore, I examine variation in sensitivity given uncertainty in tree estimation, as well as how well null congruence models align with expectations of community assembly that is completely neutral with respect to host phylogeny. I generally found that model performance increased rapidly with increasing tree sizes, suggesting lower limits on the host phylogenetic breadth needed to reliably detect phylosymbiotic signals with this approach. Furthermore, I found several notable variations in performance between congruence metrics, which translated into different inferences regarding signal detection. Overall, these findings suggest that Monte Carlo sampling across tree space can be an effective way to quantify phylosymbiotic signals and highlight key considerations for its implementation.

## Introduction

The structure and composition of host-associated microbial communities vary significantly across the tree of life. In many host taxa, this variation appears to be predominately explained by extant environmental factors that might not necessarily track the host phylogeny, such as diet or local habitat (Sullam et al., 2012, Davenport, 2016, Martinson et al., 2017, DuBose et al., 2025). In other cases, this variation mirrors the patterns of evolutionary divergence between hosts, a phenomenon known as *phylosymbiosis* (Mazel et al., 2018, Brooks et al., 2016, Lim and Bordenstein, 2020, Kohl, 2020, Perez-Lamarque et al., 2023). Patterns of phylosymbiosis can be generated by the combination of different ecological and evolutionary processes operating with varying degrees of importance (Mazel et al., 2018, Lim and Bordenstein, 2020, Kohl, 2020, Perez-Lamarque et al., 2023). In other words, the specific processes that generated any given pattern of phylosymbiosis are likely to differ between groups (Lim and Bordenstein, 2020, Kohl, 2020, Qin et al., 2023). Therefore, understanding the diversity of processes that produce emergent phylosymbiotic patterns requires quantifications of phylosymbiotic signals from across a broad range of taxa (Kohl, 2020).

To this end, there have been several approaches proposed for statistically evaluating if patterns of microbial community dissimilarity significantly parallel patterns of their host’s evolutionary relatedness. One popular approach is to transform the host phylogeny into a dissimilarity matrix and use a Mantel test (Mantel, 1967) to quantify its correlation with the corresponding microbial community dissimilarity matrix (reviewed in Lim and Bordenstein, 2020). However, the Mantel test, in addition to suffering from low power, relies on linear or monotonic associations between corresponding matrix elements, an assumption that may not always be satisfied (Perez-Lamarque et al., 2022). An alternative approach that has been more recently employed is to construct a dendrogram representing patterns of microbial community dissimilarity between hosts, and quantify the topological congruence between said dendrogram and the host’s phylogenetic tree (Lim and Bordenstein, 2020, Lin et al., 2024). Although this approach relaxes assumptions of linear correspondence between host phylogeny and microbial community dissimilarity, it provides only a single quantification that can be difficult to interpret and use to make statistically informed inferences. In response to this issue, many have used Monte Carlo simulations to generate null models of phylosymbiotic signal (Brooks et al., 2016, Trevelline et al., 2020, Grond et al., 2020). This is accomplished by either by repeatedly sampling random trees (or shuffling tip labels, which yield nearly equivalent results (Mazel et al., 2018)) and quantifying their congruence with the host phylogeny. Using this approach, phylosymbiotic signal can be statistically assessed by calculating the probability that the congruence between the host phylogeny and random trees is at least as great as the congruence between the host phylogeny and the corresponding microbial dendrogram.

The assessment of phylosymbiotic signal using this Monte Carlo-based framework fundamentally relies on quantification of tree congruence, a task that can be approached using a variety of existing metrics. The Robinson-Foulds (RF) distance, which has historically been the most widely used metric, measures the number of splits that are present in one tree but not the other (Robinson and Foulds, 1981). Although widely used, this calculation has been criticized as too conservative because it does not account for similar but not identical splits (Smith, 2022). Therefore, a variety of Robinson-Foulds inspired approaches for addressing this issue have been developed, among the more notable of which are the information-corrected Robinson-Foulds distance (ICRF) (Smith, 2020a), Nye similarity (NS) (Nye et al., 2006), and the Jaccard-Robinson-Foulds distance (JRF) (Böcker et al., 2013). The matching split distance (MSD), another popular metric, takes a different perspective on this issue by finding the matching of splits between trees that minimize dissimilarity (Bogdanowicz and Giaro, 2012). More recently, information-theoretic approaches have been introduced to weight splits based on their information content, which has significantly improved quantification of tree congruence (Smith, 2020a). In addition to the previously described ICRF, this has led to the development of new information-theoretic tree congruence metrics, such as matching split information distance (MSID), mutual clustering information (MCI), and shared phylogenetic information (SPI) (Smith, 2020a).

The previously described metrics represent popular choices across the various fields that frequently evaluate tree congruence. However, there is reason to suspect their utility for assessing phylosymbiotic signals may vary due to how they emphasize and penalize different features of congruence. For example, the Robinson-Foulds derivatives emphasize overall tree structural similarity more than individual splits, while the information-theoretic approaches place less emphasis on uninformative differences. In other words, Robinson-Foulds derivatives will define trees as similar because most splits are close matches, and information-theoretic approaches will define trees as similar because they share the most informative splits. In practice, it is unclear if these differences translate into different inferences regarding statistical support for phylosymbiotic associations. In addition to uncertainty regarding the effects of congruence metric, how broad of host phylogenetic coverage is needed to detect a phylosymbiotic signal using the previously described Monte Carlo-based approach is also unclear. In other words, it is important to know if the attained breadth of host phylogenetic coverage is sufficient for detecting a phylosymbiotic signal, particuarly given the inherent noise associated with variation in host-associated microbial communities.

To this end, here I examined the variation in performance associated with the aforementioned null model approach when used with different extents of host phylogenetic breadth and congruence metrics. Through simulations under null and alternative hypotheses, I found that null model performance generally increased rapidly with increasing tree size. In this, good power and type 1 error (false positive) rates were typically achieved for trees of between 10-20 leaves. However, I also found several minor variations in performance between congruence metrics, which translated into their frequent disagreement regarding the significance of phylosymbiotic associations. Taken together, these findings provide useful considerations for practitioners by illustrating the conditions in which this null model approach is likely sufficient or insufficient for quantifying patterns of phylosymbiosis.

## Methods

### Monte Carlo algorithm for generating null models of phylosymbiosis

Let the host phylogeny be *T*_*h*_ and the associated microbial community dendrogram be *T*_*c*_, each with *l* leaves. The observed congruence is then *O* = *C* (*T*_*h*_, *T*_*c*_ ), where *C* represents a previously described congruence metric. To generate the null model, *n* unrooted binary trees with *l* leaves are randomly sampled with replacement from a uniform tree distribution (all topologies are equally likely), denoted as defined as 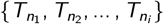. The null congruence distribution is then defined as *N* = {*C*_1_, *C*_2_, …, *C*_*n*_}, where 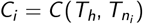. In other words, the null congruence distribution is the congruence vector between the host tree and each of the randomly drawn trees. This distribution can then be used to test the null hypothesis *H*_0_ : *O* ≤ *N*, against the alternative hypothesis *H*_*A*_ : *O > N*. These hypotheses can then be assessed by calculating the proportion of the null distribution that is greater than or equal to the observed congruence (since this is a one-sided test):

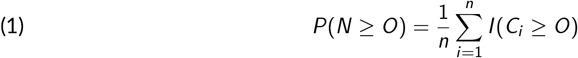

where *I* is the indicator function, equal to 1 if the inequality is satisfied and 0 otherwise. Note that for some metrics, greater congruence is associated with a lower value, in which case the direction of the inequality is reversed. A graphical representation of this processes is depicted in Figure 1. For convenience, I hereafter refer to this method as a random tree congruence test. For methodological evaluations described in this text, as well as promoting use of this approach, I have implemented this algorithm in *manticore* (Monte cArlo simulaioN algoriThm for assessIng tree COngRuencE), which is an open-source R library that provides a simple interface for its utilization. Within *manticore*, the *rtopology* function from the *ape* R library (v5.7.1) is used to sample random trees (Paradis and Schliep, 2019) and various functions from the *TreeDist* R library (v2.7.0) are used for quantifying congruence (Smith, 2020b).

**Figure 1.**
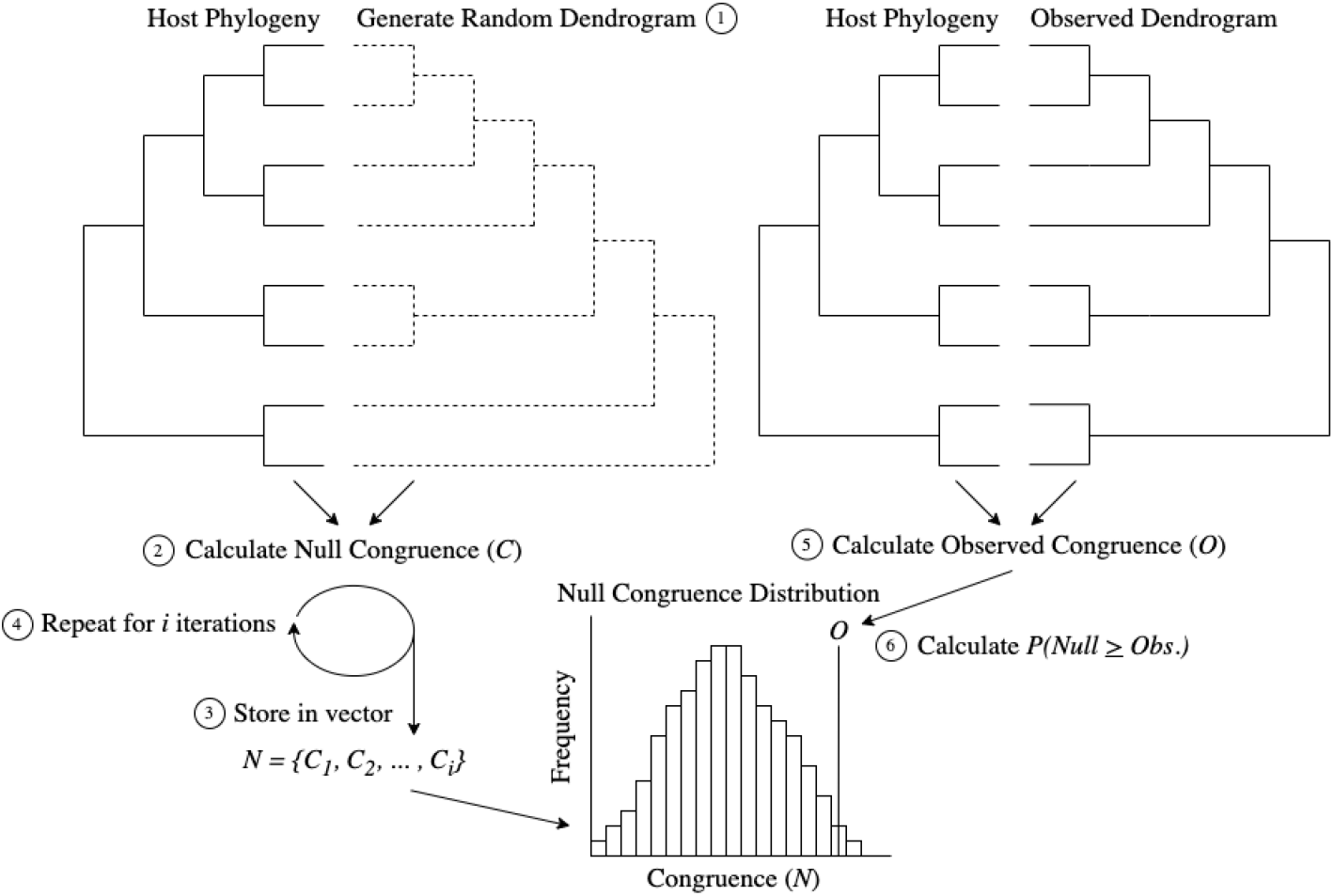
A graphical depiction of the Monte Carlo algorithm for generating null congruence distributions. The processes starts by (1) sampling a random dendrogram and (2) calculating how congruent it is to the host phylogeny. This congruence value (denoted *C* ) is stored in a vector, and (4) steps 1-3 are repeated for *i* iterations. The resulting vector then serves as the null distribution that will be used for hypothesis testing. (5) Calculate the observed congruence between the host phylogeny and the corresponding dendrogram (denoted *O*). Then (6) use the previously described null distribution to calculate the probability that randomly generated trees that showed greater than or equal to congruence to the host phylogeny relative to the observed congruence.

### Simulations under *H*_0_ and *H*_*A*_

To evaluate the type 1 error (false positive) rate associated with each metric when used in random tree congruence testing, I first simulated 1000 pairs of independently randomly sampled host phylogenies and microbial community dendrograms (under *H*_0_) for trees with 5, 10, 20, 40, 60, 80, and 100 leaves. For each pair, I conducted a random tree congruence test using each congruence metric with 1000 iterations and recorded *P*(*Null* ≥ *Observed* ) values.

To evaluate the power (true positive rates) associated with each metric when used in random tree congruence testing, I simulated variation in microbial community dissimilarities using a subtree pruning and regrafting (SPR) approach to mutate host phylogenies (under *H*_*A*_). Specifically, this process first generates a host phylogeny. To generate associated microbial community dendrograms, the host phylogeny is then subject to a number of subtree prunings and regraftings, which I varied to represent portions of the total leaves that ranged from 0 to 0.5, by 0.1. This process recapitulates what could be expected if factors that do not necessarily track host’s phylogeny (e.g., diet, local environment) also play a role in shaping their associated microbial communities. In other words, more subtree pruning and regrafting introduces more noise in the phylosymbiotic association. For each previously described tree size and congruence metric, I conducted 50 replicate simulations, and calculated the proportion of significant tests as a function of the number of subtree prunes and regrafts. I used the *SPR* function from the *phangorn* R library (v.2.11.1) for all subtree pruning and regrafting (Schliep, 2011).

### Evaluating true and false positive rates

To evaluate the power and false positive rates associated with each metric and aforementioned tree size, I used the previously described simulations to construct receiver operating characteristic (ROC) curves. ROC curves describe how well a statistical test distinguishes between the null and alternative hypotheses based on the *P*(*Null* ≥ *Observed* ) values attained from both simulations. To describe performance more quantitatively, I calculated the area under each ROC curve (AUC). Here, values closer to 1 indicate that the distribution of *P*(*Null* ≥ *Observed* ) attained from the *H*_*A*_ and *H*_0_ simulations are more distinguishable, while values closer to 0.5 indicate they are less distinguishable. I performed all ROC curve construction and AUC calculations using the *SciPy* Python library (v.1.13.1) (Virtanen et al., 2020).

It is important to note that ROC curve construction utilized all *H*_*A*_ simulations, which span a gradient of noise levels. This provides a more general picture of how this test can be expected to perform in practice, but it does not provide intuition about how power may vary with increased noise, or the interaction of this function with tree size. Therefore, I also constructed curves that examine power, which is the proportion of significant results at *α* = 0.05 under *H*_*A*_ simulations, as a function of the extent of subtree pruning and regrafting. To facilitate comparison across tree sizes, I divided the number of subtree prunes and regrafts by tree size. Similarly, I also constructed curves that examine type 1 error (false positive) rates as a function of tree size. Specifically, I calculated the proportion of simulations under *H*_0_ that were greater than or equal to *α* = 0.05, where the expected type 1 error rate for a well calibrated test should be approximately 0.05 for said *α*.

### Sensitivity analyses

The standard random tree congruence test treats both the host phylogenetic tree and microbial community dendrograms as fixed objects. However, there is typically error associated with estimation of both trees. One approach to understanding how this error may impact biological inferences would be to propagate error from, for example, phylogenetic or microbial community bootstrapping. However, since there are many approaches researchers use to construct said trees, investigations regarding any one or subset of these approaches may not be generalizable. Therefore, I instead focus on sensitivity analyses that generally perturb tree topologies, which allows for assessment of how likely inferences are to change given uncertainty in tree estimation. Here, instead of setting the congruence between a host tree microbial dendrogram as a fixed value, I generated distributions of *k* possible observed congruence values *O*_*p*_ = {*O*_1_, *O*_2_ … *O*_*k*_ } by implementing increasing degrees of subtree pruning and regrafting. Given that each *O*_*p*_ is equally likely (due to uniform sampling), a Monte Carlo approximation can be used to calculate an integrated probability of observing null congruence values greater than or equal to those of the congruence distribution:

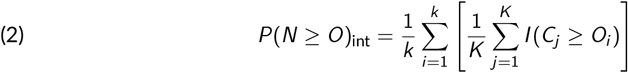

By increasing the degrees of subtree pruning and regrafting (relative to the number of tree leaves), one can identify at what level of uncertainty (if any) results are not robust (where *P*(*N* ≥ *O*)_int_ ≥ *α*). To examine variation in sensitivity to uncertainty across metrics, I first sampled host trees and issued minor perturbations of N SPRs = *n*_leaves_·0.05 to generate microbial dendrograms with a small degree of noise. For each pair of trees, constructed null congruence models using 1000 iterations and calculated *P*(*N* ≥ *O*)_int_ as previously described. For simplicity, I conducted 20 replicate simulations for each metric and each tree size of 20, 60, and 100 leaves. Within each of these, I generated congruence distributions for each 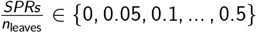 to evaluate where *P*(*N* ≥ *O*)_int_ ≥ *α*.

### Null congruence model comparison to totally neutral microbial assembly

Vellend’s foundational synthesis in community ecology describes the four fundamental processes that drive the dynamics of ecological communities as analogous to those that drive the dynamics of alleles within populations. That is, diversification (mutation) and dispersal (gene flow) can introduce taxa into a community, and ecological selection and drift then shape their frequencies (Vellend, 2010). Patterns of phylosymbiosis can arise from any combination of these processes working in tandem (Lim and Bordenstein, 2020). The most discussed is when host-associated microbial communities are selectively assembled with respect to their host’s phylogeny. However, phylosymbiotic patterns may also be generated as a product of neutral assembly and dispersal limitation, where microbial taxa within a given host’s niche are less likely to interact with hosts that occupy more distant niches (Lim and Bordenstein, 2020). While this list could go on, the general point is that any combination of ecological processes can give rise to phylosymbiotic signals.

Since the approach proposed here is meant to serve as a null model, the primary ecological process that a useful null model should mirror is microbial community assembly that is completely neutral with respect to host phylogeny. Therefore, I investigated how well null congruence models correspond to totally neutral assembly with respect to host phylogeny. For each metric and tree size, I simulated 100 pairs of trees and constructed random tree null congruence distributions with 1000 iterations. I also simulated congruence distributions from neutrally assembled microbial communities. Let *M* be the number of microbial taxa, *N* be the community size (per host), and *H* be the number of hosts. Each microbial taxon is equally likely to colonize each host. Therefore, letting *p* = (*p*_1_, *p*_2_, … *p*_*S*_ ) be the vector of probabilities that each microbial taxon *p*_*i*_ colonizes a given host, where each 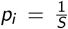, the microbial community associated with each host is assembled as *X*_*h*_ ∼ Multinomial(*N, p*). For computational convenience, I ran all simulations with *M* = 100 and *N* = 1000. I then calculated the pairwise Bray-Curtis distance between hosts, and created microbial community dendrograms using hierarchical clustering (UP-GMA). For Bray-Curtis calculations I used the *vegdist* function from the *vegan* R library (v.2.6-4) (Oksanen et al., 2022) and for hierarchical clustering I used the base R *hclust* function (v.4.2.0) (R Core Team, 2022).

For each simulation iteration, I then examined how well random tree null congruence distributions corresponded with congruence distributions generated from neutral assembly. Let *F*_random tree_(*x* ) and *F*_neutral_(*x* ) be the empirical cumulative density functions for random tree null congruence distributions and neutral assembly congruence distributions, respectively. The normalized difference between distributions relativized to their maximal total difference is

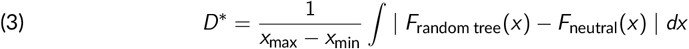

Here, *D*^∗^ ∈ [0, 1], where *D*^∗^ = 0 indicates distributions are identical and *D*^∗^ = 1 indicates distributions are completely non-overlapping. For each metric, I evaluated *D*^∗^ as a function of tree size by fitting a saturating function *f* (*x* ) = *a*(1 − *e*^−*bx*^ ) to simulation results (due to visible saturation in some metrics).

## Results

### Determining sufficient simulation effort

Due to their random nature, the number of simulations to conduct is an important feature to consider when making inferences based on Monte Carlo-based null models. Previous studies have handled this issue by performing an excessively large number of simulations. However, this approach may be too computationally demanding for some studies, particularly those that might wish to conduct scans across many sets of trees (e.g., Pollock et al., 2018). Fortunately, because Monte Carlo simulation is binomial sampling, known properties of the binomial distribution can be used to assess whether simulation efforts were sufficient. Let *p* = *P*(*Null* ≥ *Observed* ) represent the true p-value one would obtain from exact probability calculations or infinite Monte Carlo simulations. Using finite sampling, *p* is estimated by conducting *i* replicate simulations and calculating the proportion 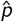 of simulations where the null statistic is greater than or equal to the observed. Because this is equivalent to binomial sampling, the standard error of 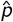 is:

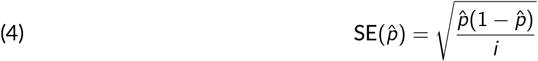

This allows one to assess uncertainty in the estimated p-value without conducting excessive simulations. For example, if *ε* is the desired standard error threshold, the number of simulations conducted is sufficient if

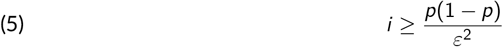

Furthermore, although less commonly applied, a 95% confidence interval can be estimated as

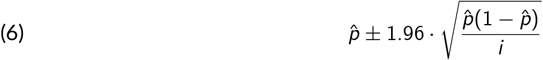

Practitioners can use this framework to determine the number of simulations necessary to make confident inferences. For example, if lower bound *<* upper bound *< α*, one can confidently reject the null. Likewise, if lower bound *< α <* upper bound, the simulation is too uncertain and more simulations should be conducted. One can also use this logic to specify *ε* and therefore solve for the number of simulations needed. Intuitively, the margin of error should be less than the distance between 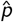 and *α*. This can be used to redefine Equation 5 in terms of *α*:

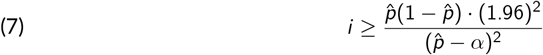

While this framework is useful for practitioners to consider, it also contextualizes uncertainty in the remaining simulation study. For example, since the maximum standard error occurs when 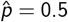, the approximate margin of error for simulations with 1000 iterations is ≈ 0.03. Therefore, for *α* = 0.05 as long as 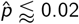 or 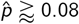, inferences are expected to be stable.

### Test performance rapidly increases with tree size and shows minor variation by congruence metric

To assess how well random tree congruence testing discriminates between null and alternative simulations based on the *P*(*Null* ≥ *Observed* ) attained from each, I constructed ROC curves that show the true positive rate as function of the false positive rate. Here, curves that rapidly approach a true positive rate of 1 indicate that the test is correctly discriminating true positives from false positives. Visual inspection of these curves across congruence metrics generally show this pattern is typically achieved for trees with 20 leaves or greater (Figure 2). Conversely, the test was less able to discriminate true from false positives for trees with 10 leaves, and even less able for trees with 5 leaves. The aforementioned pattern was general across congruence metrics, with the exception of ICRF. Here, use of ICRF resulted in decreased discriminatory power from across tree sizes (Figure 2).

**Figure 2.**
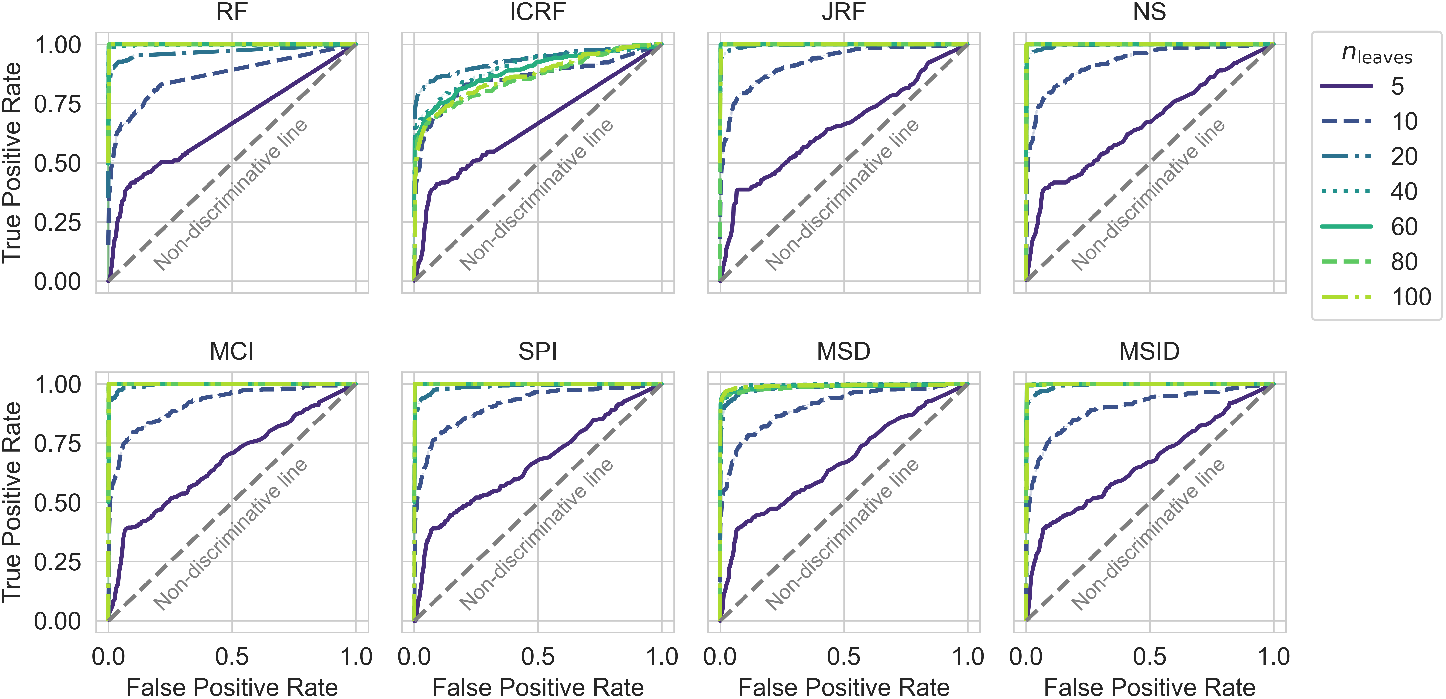
Receiver operating characteristic (ROC) curves showing the random tree congruence test’s ability to discriminate between true and false positives. Each panel corresponds to a different congruence metric and shows the true positive rate (y-axis) as a function of false positive rate (x-axis). Different colored and patterned lines represent the ROC curve for different tree sizes. The gray y=x line in each panel represents the non-discriminative line, which is the expected function for a test that is completely unable to discriminate between true and false positives.

To more quantitatively examine the previously described trends, I plotted the area under each ROC curve as a function of tree size. This showed that for all congruence metrics besides ICRF, values of AUC ≈ 1 were achieved for trees with 20 leaves and greater (Figure 3). Likewise, the test showed significantly reduced discriminative ability for trees with 10 leaves, which was further reduced to near the non-discriminative line for trees with 5 leaves (Figure 3). Consistent with my visual account of the ROC curves, use of ICRF resulted in decreased discriminative ability across tree sizes. Although there was a significant increase in area under ROC curve from trees with 5 leaves to 20 leaves, use of this metric never resulted in AUC ≈ 1 for any tree size (Figure 3).

**Figure 3.**
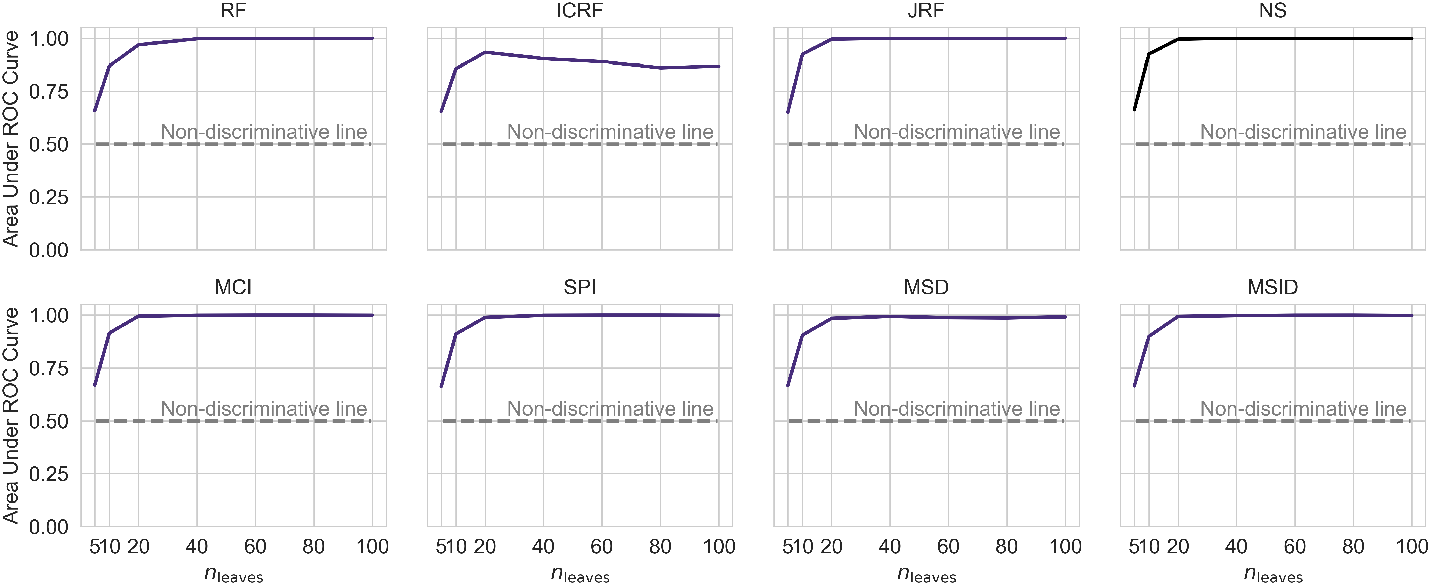
Ability to discriminate between true and false positives rapidly increases with tree size. Each panel corresponds to a different congruence metric and shows the area under the ROC curve (y-axis) as a function of tree size (x-axis). Here, values closer to 1 indicate greater ability to discriminate between true and false positives, while values closer to 0.5 represent total inability to discriminate between true and false positives (denoted as the gray dashed line in each panel).

### Declines in power associated with increasing noise are generally recovered in larger tree sizes but vary by congruence metric

The previously described analyses considered the test’s general ability to discriminate between true and false positives, but did not consider how true positive detection varies with increasing noise. To evaluate this property, I plotted power as a function of noise level. Here, power is the proportion of significant results under *H*_*A*_ simulations at *α* = 0.05, and noise level is the number of subtree prunes and regraphs divided by the number of leaves (to facilitate cross comparison between tree sizes). Examining said curves showed several general patterns. First, consistent with the previously described analyses, virtually no trees with only 5 leaves were detected as significant (Figure 4). However, the test did show some power to detect true positives for trees with 10 leaves at lower levels of noise. The noise level at which this curve drops below a power of 0.8 (a typical standard) was generally between 0.2 and 0.3. Use of RF and ICRF reduced this noise level to between 0.1 and 0.2 (Figure 4), indicating less power.

**Figure 4.**
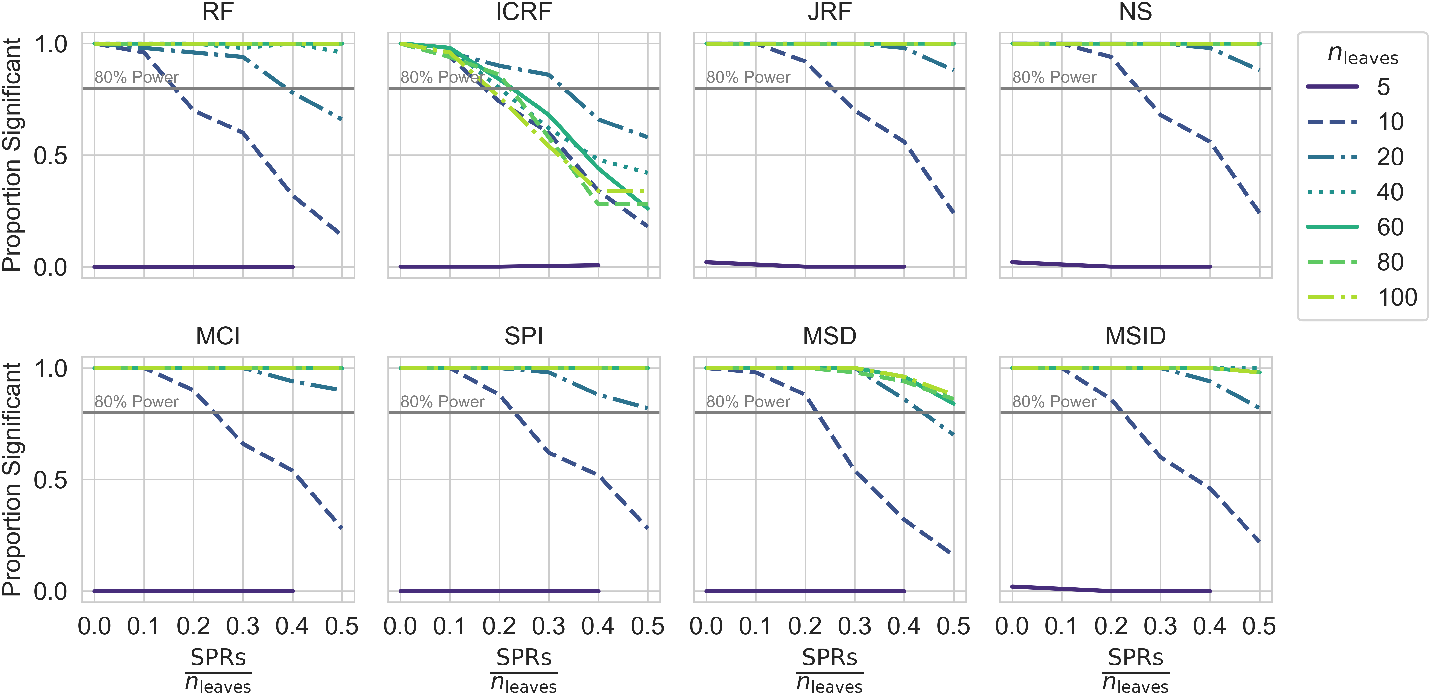
Variation in the relationship between power and noise level by tree size and congruence metric. Each panel corresponds to a different congruence metric and shows the proportion of significant results under *H*_*A*_ simulations (power) (y-axis) as a function the number of subtree prunes and regrafts divided by the number of leaves (noise level) (x-axis). Each panel also has a gray line that depicts a power level of 0.8, which is a standard benchmark for concluding sufficient power.

Increasing tree size to 20 leaves recovered a significant degree of power across congruence metrics. Notably, use of JRF, NS, MCI, SPI, and MSID resulted in acceptable degrees of power (≥ 0.8) across the full span of noise tested, although there was a decline in power at higher levels (Figure 4). Alternatively, use of MSD did confer greater power across the noise spectrum, but power levels did drop below acceptable at high levels of noise (between 0.4 and 0.5) (Figure 4). Use of RF and ICRF still resulted in significantly reduced power, which dropped below acceptable levels at noise levels of between 0.3 and 0.4 (Figure 4). Finally, further increases to tree size (*n*_leaves_ ≥ 40) generally recovered acceptable power across the noise spectrum (note that although use of MSD still showed declines with higher levels of noise, power did not drop below 0.8). The power associated with use of ICRF proved to be an exception to this pattern, which generally dropped below the acceptable level of 0.8 at noise levels of ≈ 0.2 for even larger tree sizes (Figure 4).

### Variation in Type 1 error rate by congruence metric and tree size

Similar to the analyses presented in previous section, it is useful to decompose ROC curves to examine if and how type 1 error (false positive) rates vary across congruence metrics and tree sizes. Therefore, I visualized the proportion of simulations under *H*_0_ that were significant (*α* = 0.05). A well calibrated test should show that said rate is approximately equal to 0.05, which is indeed the general case across congruence metrics and tree sizes (Figure 5). However, there are two notable deviations from this trend. First, the test did not reliably identify significant signals of congruence for trees with 5 leaves, regardless of congruence metric (Figure 5). This is unsurprising given the shapes of the ROC curves. Second, use of RF conferred significantly lower type 1 error rates than expected across tree sizes, indicating its conservative nature (Figure 5).

**Figure 5.**
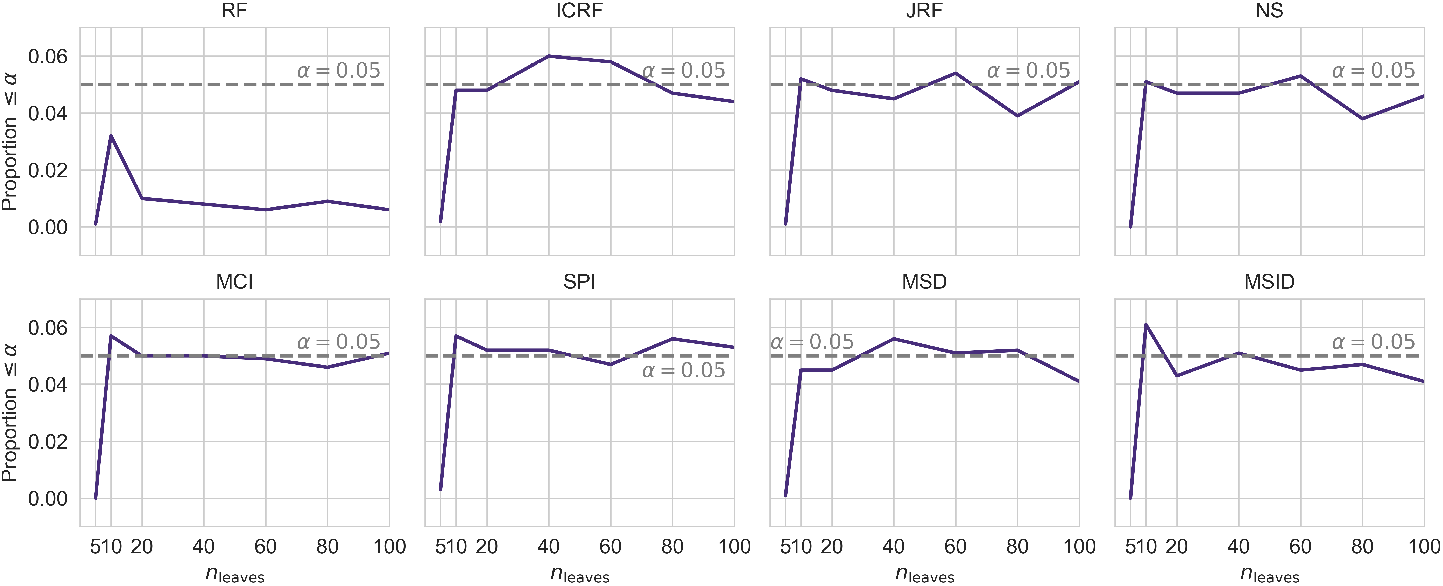
Variation in type 1 error rates associated with different tree sizes and congruence metrics. Each panel corresponds to a different congruence metric and shows the proportion of simulations under *H*_0_ that were significant at *α* = 0.05 (y-axis) as a function of tree size (x-axis). The gray in each panel depicts the expected type 1 error rate for the corresponding *α*.

### Variation in sensitivity to uncertainty in tree estimation by metric and tree size

The previous analyses treat host and microbial community trees as fixed objects. However, there is typically a degree of uncertainty associated with tree estimation. To consider how much uncertainty can be handled before inferences change, I generated congruence distributions by using increasing degrees of subtree pruning and regrafting to perturb host and microbial community trees. I then evaluated these congruence distributions, rather than a fixed congruence value, against the null and examined how the integrated *P*(*Null* ≥ *Obs*.) changed with increasing degrees of perturbation. This showed use of JRF, NS, MCI, and SPI generally conferred the most robustness to uncertainty, though trees of *n*_leaves_ = 20 offered less robust inferences at moderate degrees of uncertainty (Figure 6). Use of RF also showed reasonable robustness to uncertainty. However, this was primarily the case for larger trees, as inferences for *n*_leaves_ = 20 were less robust at lower degrees of uncertainty (Figure 6). Similarly, use of MSD and MSID did not confer as much robustness for larger trees, but trees of all sizes exhibited similar degrees of robustness at lower-middle levels of uncertainty (Figure 6). Finally, use of ICRF conferred the least robustness to uncertainty, as inferences (at *α* = 0.05) changed for trees of all sizes at low degrees of uncertainty (Figure 6).

**Figure 6.**
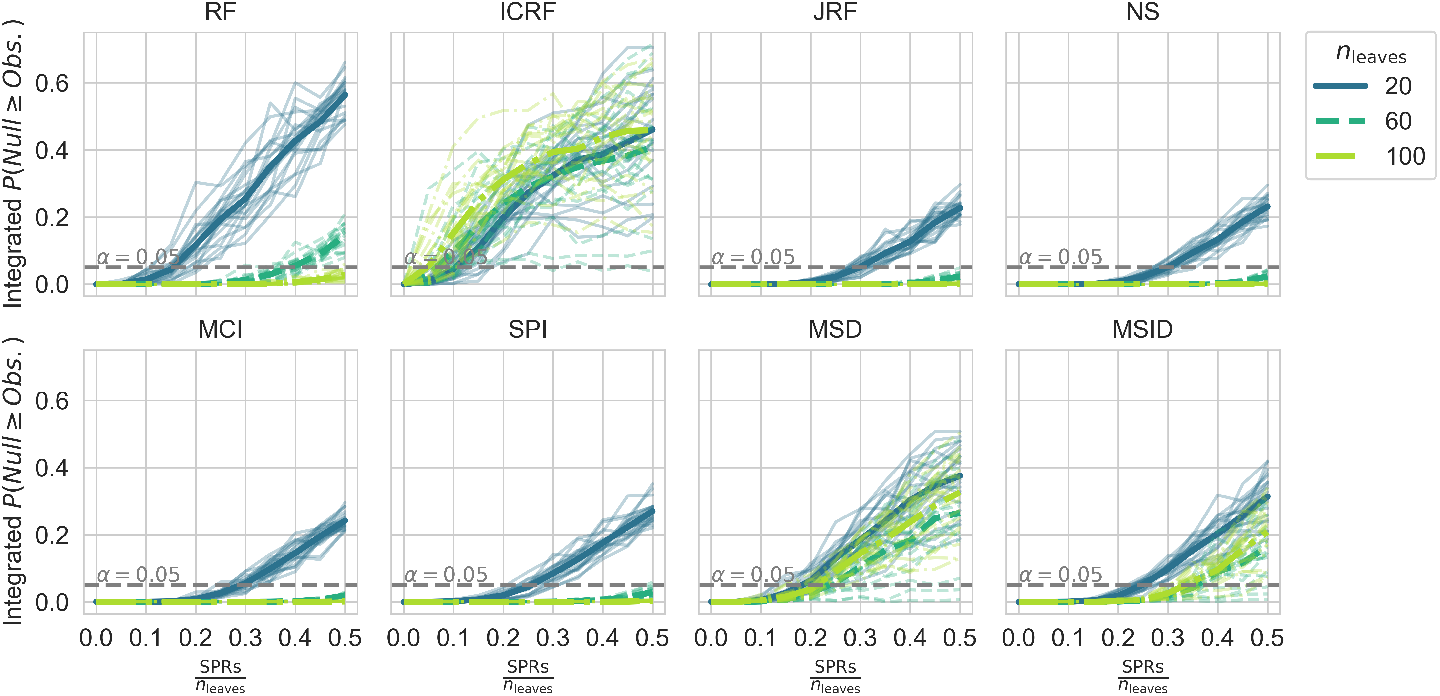
Variation in sensitivity to uncertainty associated with different tree sizes and congruence metrics. Each panel corresponds to a different congruence metric and shows the integrated *P*(*Null* ≥ *Obs*.) as a function of 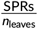 (issued degree of uncertainty). Each fainter line represents a replicate simulation run, and each darker line represents the average across simulations. The dashed gray line indicates *α* = 0.05, which is the typical threshold for changing statistical inference.

### Similarity between random tree congruence models and neutral community assembly models varies by congruence metric and tree size

The only ecological process that cannot produce patterns of phylosymbiosis is totally neutral community assembly, where all microbial taxa have equal probabilities/rates of colonizing a set of possible host taxa. Therefore, a useful null congruence model should show general agreement with the congruence distribution from neutrally assembled host-associated microbial communities. To evaluate this agreement, I quantified the (normalized) integrated difference between random tree congruence distributions and congruence distributions from neutrally assembled communities and examined how this quantity changes with tree size. This showed significant variation between metrics. Overall, RF showed a significant degree of similarity (*D*^∗^ *<<* 0.2) with neutral assembly congruence distributions, which did not significantly vary across tree sizes (Figure 7). MCI and SPI both showed reasonable similarity with neutral assembly congruence distributions for smaller trees, but this did increase to *D*^∗^ ≈ 0.2 in larger trees (Figure 7). JRF and NS showed similar patterns but with a significant degree of heteroscedasticity, where the variation across simulations increased as tree size increased (Figure 7). MSD and MSID showed even greater increases in divergence from neutral assembly congruence distributions with increasing tree size, exceeding *D*^∗^ ⪆ 0.2 between trees with 20-30 leaves (Figure 7). Finally, ICRF showed a rapid increase to *D*^∗^ ≈ 0.4 at trees with 20 leaves before plateauing at *D*^∗^ ≈ 0.5, indicating significant deviation from neutral assembly congruence distributions (Figure 7).

**Figure 7.**
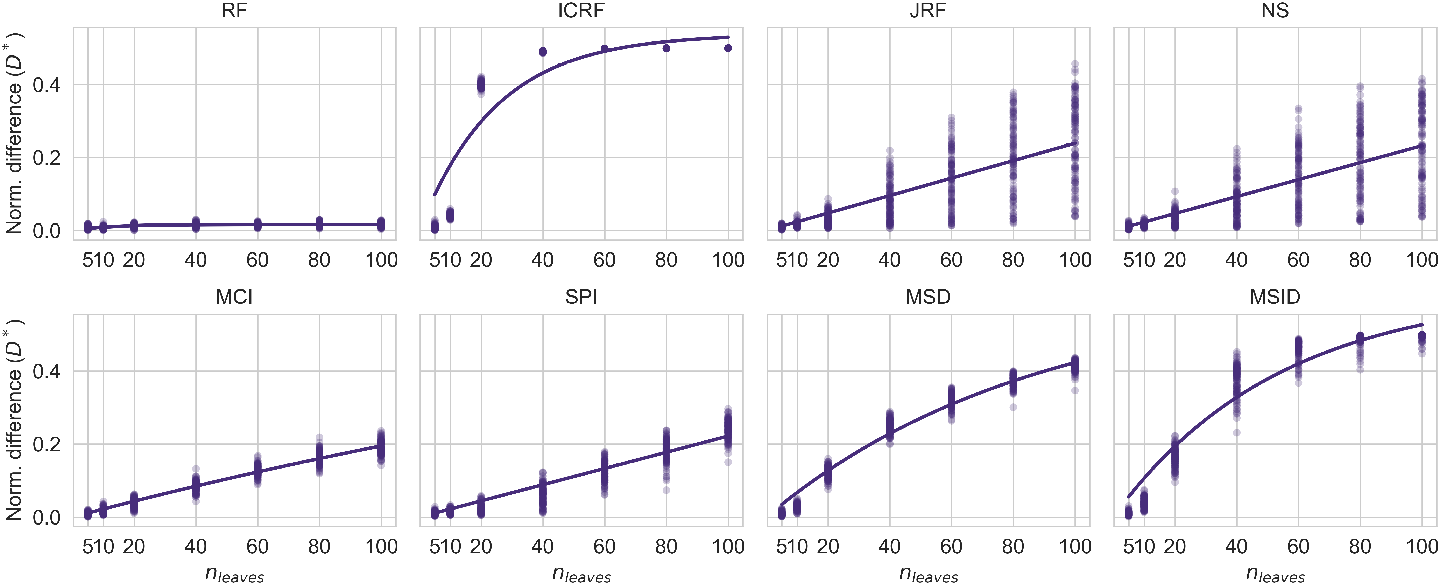
Variation in similarity between random tree congruence models and neutral community assembly models rates associated with different tree sizes and congruence metrics. Each panel corresponds to a different congruence metric and shows changes in normalized differences *D*^∗^ (Equation 3) as a function of tree size. Points represent individual simulations, and lines represent the saturating function fit to simulated data for trend visualization.

### Use of different congruence metrics generally lead to the same inference regarding the detection of a phylosymbiotic signal

Now that the power, type 1 error, and agreement with neutral assembly associated with different congruence metrics have been described, it is important to ask whether or not these differences translate into meaningfully different inferences regarding whether a phylosymbiotic relationship is statistically supported. To answer this question, I used the previously described *H*_*A*_ and *H*_0_ simulations to calculate the proportion of times each pair of metrics agreed on whether a phylosymbiotic relationship was statistically supported (at *α* = 0.05). Overall, this showed that for both *H*_*A*_ (Figure 8A) and *H*_0_ (Figure 8B) simulations different metrics tended to result in the same statistical inference. Specifically, most metrics agreed between 93 − 100% of the time. Unsurprisingly, the notable exception to this was ICRF, which showed agreement approximately 80% of the time with all other metrics (Figure 8A). However, higher levels of agreement were achieved in *H*_0_ simulations (Figure 8B), which is consistent with the previous differences in power and type 1 error rates associated with ICRF.

**Figure 8.**
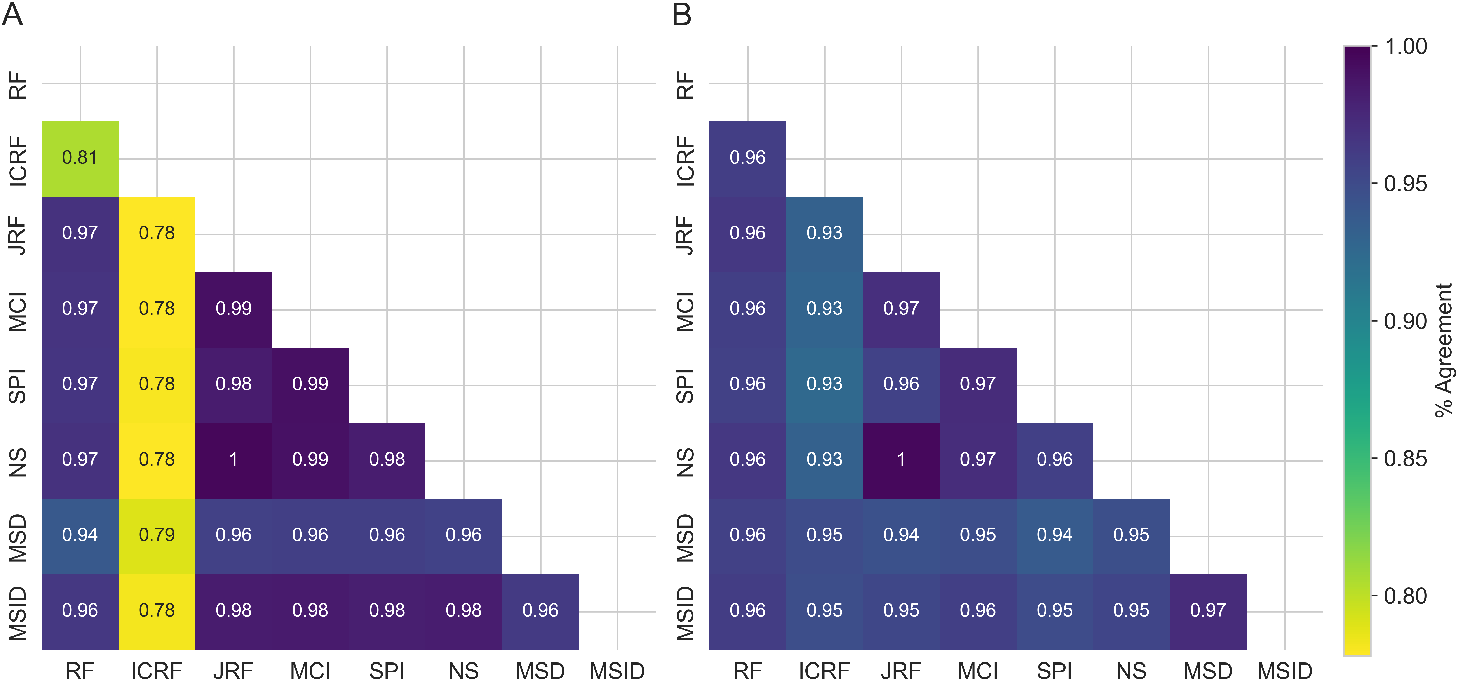
Use of different congruence metrics tend to result in consistent inferences regarding the statistical support of phylosymbiotic relationships. Panels show heatmaps that depicts the proportion of simulations where the corresponding metrics agreed on the statistical support for a phylosymbiotic relationship (at *α* = 0.05) across A) *H*_*A*_ simulations and B) *H*_0_ simulations. The number in each cell indicates the proportion of agreements for the corresponding metrics.

## Discussion

Understanding patterns of phylosymbiosis relies on our ability to detect and quantify said patterns. Here, I examined the suitability and nuances of a previously employed Monte Carlo-based null model approach for accomplishing this task (Brooks et al., 2016, Mazel et al., 2018, Trevelline et al., 2020, Grond et al., 2020). Through null and alternative model simulations, I found that this approach generally showed adequate power and false discovery rates for trees of sufficient size (≥ 10 leaves) (Figures 2, 3, 4, and 5). However, there was significant variation in how sensitive null models constructed with different congruence metrics were to uncertainty in tree estimation (Figure 6). Likewise, there was significant variation in how well null models constructed with different congruence metrics corresponded to null models of neutral community assembly with respect to host phylogeny (Figure 7). Nonetheless, use of different congruence metrics generally lent consistent inferences with regards to the presences/absence of a phylosymbiotic symbol (Figure 8). Use of ICRF represents a notable exception to these generalizations, which was associated with decreased discriminatory power between true and false positives, increased sensitivity to uncertainty, and significant deviation from models of neutral assembly. These features translated into lower levels of overall agreement, particularly regarding support for true phylosymbiotic signals, with other metrics.

### Null model assumptions, limitations, and justifications

Patterns of phylosymbiosis can be generated by any combination of fundamental ecological processes. For example, differences in microbial dispersal rates to different hosts (either due to differences in transmission rates or differences in density in the environment/regional pool), differences in relative fitness within different hosts, and differences in co-evolutionary histories, are all process that can generate phylosymbiotic patterns (Lim and Bordenstein, 2020, Kohl, 2020). To disentangle these complexities, one must first test for the presence of a phylosymbiotic signal. While any combination of the previously described processes could be involved in the production of a phylosymbiotic relationship, microbial community assembly that is completely neutral (microbial taxa have equal probabilities of arriving in any given host, as well as equal fitness within each host) cannot produce patterns of phylosymbiosis (Brooks et al., 2016). Therefore, null models of phylosymbiosis fundamentally assume a lack of differences in microbial dispersal rates to different hosts, a lack of relative fitness within different hosts, and no co-evolutionary history. It is notable that only the use of RF generated null models that closely aligned with neutral expectations, regardless of tree size (Figure 7). Use of MSD and SPI somewhat aligned with neutral expectations, though with a modest degree of deviation with increasing tree size. However, use of all other metrics showed more significant increases in deviation from neutral expectations with increasing tree size (Figure 7).

Although assuming away all ecological processes besides ecological drift can provide a null exception, it offers no help in resolving the processes that generated any observed deviations from said null. Explicitly modeling ecological and evolutionary processes would be more useful for establishing mechanistic bases of phylosymbiotic patterns. However, such models fundamentally rely on stronger assumptions that may not always be generalizable (Perez-Lamarque et al., 2022). To resolve such issues, experimental tests are needed that control different processes while manipulating others (Brooks et al., 2016). The results from such experiments could then be used to inform more process-oriented models, which would ultimately lend the strongest inferences. Nonetheless, this null model approach can be applied to natural data and can help researchers initially examine the presence of phylosymbiotic patterns.

Constructing null models by sampling from a uniform tree distributions is another assumption that warrants further discussion. Alternative approaches would include generating null trees from an evolutionary process model, such as a Birth-Death diversification model or multivariate phylogenetic models of trait evolution (Morlon et al., 2010, Perez-Lamarque et al., 2023). Such approaches may produce trees with topologies that more closely resemble empirical trees relative to sampling from a uniform distribution. However, the shapes of trees are influenced by the nature of the evolutionary processes that generated diversification patterns (Colijn and Plazzotta, 2018). This null model approach is intended to represent the absence of any evolutionary relationship between hosts and their microbial communities. Therefore, sampling from a uniform tree distribution ensures that every tree is equally likely, thus ensuring a lack of shared structure due to shared diversification processes. Conversely, utilizing for example a Birth-Death model would introduce structured correlations that might inadvertently resemble non-random diversification, potentially biasing null distributions. Therefore, sampling from a uniform distribution provides a more unbiased test the null hypothesis that the observed congruence is due to chance (pure stochasticity).

### Practical recommendations

Overall, I found notable difference in null model performance associated with different congruence metrics that may be useful to consider. Notably, use of ICRF conferred decreased power which led to decreased ability to distinguish between true and false positives. Furthermore, use of ICRF provided less robust inferences given uncertainty in tree estimation, and showed significant deviations from neutral assembly expectations. Taken together, these findings suggest use of ICRF may not be sufficient for reliably detecting phylosymbiotic signals. Alternatively, use of JFR, NS, MCI, and SPI all showed good power, error rates, and robustness given tree uncertainty. However, MCI and SPI showed greater consistency with expectations from totally neutral assembly. Use of MSD and MSID performed similarly in terms of power, error rate, but their sensitivity to uncertainty and significant deviations from neutral assembly expectations make their use less defensible. Finally, use of RF conferred somewhat less (but still decent) power in smaller trees, but showed significantly reduced degrees of type 1 error, illustrating the conservative nature of RF. Use of RF also showed less robustness to tree estimation uncertainty for smaller trees, but decent robustness in larger trees. Despite its somewhat weaker statistical performance relative to JFR, NS, MCI, and SPI, it showed negligible differences from expectations of totally neutral assembly, suggesting it may serve as a more principled approach for null model generation.

Overall, these findings suggest use of RF may provide a more conservative yet principled inference of phylosymbiotic signal, but its use should be limited to larger trees or smaller trees with lower degrees of uncertainty. Given higher degrees of uncertainty or desire for a less conservative estimate, use MCI and SPI would be more optimal.

## Data, script, code, and supplementary information availability

All code written for implementing the algorithm described here has been packed as an R library called *manticore*, which is available on GitHub athttps://github.com/gabe-dubose/ manticore and archived with Zenodo at https://doi.org/10.5281/zenodo.14828441 (DuBose, 2025). All scripts written for the benchmarks and evaluations presented in this manuscript are present in the same GitHub repository but under the “evaluation” directory (https://github.com/gabe-dubose/manticore/tree/main/evaluation).

